# Beyond the MEP Pathway: a novel kinase required for prenol utilization by malaria parasites

**DOI:** 10.1101/2023.07.17.549440

**Authors:** Marcell Crispim, Ignasi Bofill Verdaguer, Agustín Hernández, Thales Kronenberger, Àngel Fenollar, María Pía Alberione, Miriam Ramirez, Alejandro Miguel Katzin, Luis Izquierdo

## Abstract

A promising treatment for malaria is a combination of fosmidomycin and clindamycin. Both compounds inhibit the methylerythritol 4-phosphate (MEP) pathway, the parasitic source of farnesyl and geranylgeranyl pyrophosphate (FPP and GGPP, respectively). Both FPP and GGPP are crucial for the biosynthesis of several essential metabolites such as ubiquinone and dolichol, as well as for protein prenylation. Dietary prenols, such as farnesol (FOH) and geranylgeraniol (GGOH), can rescue parasites from MEP inhibitors, suggesting the existence of a missing pathway for prenol salvage via phosphorylation, by competition. In this study, we identified a gene in the genome of *P. falciparum*, encoding a transmembrane prenol kinase (PolK) involved in the salvage of FOH and GGOH. The enzyme was expressed in *Saccharomyces cerevisiae*, and its FOH/GGOH kinase activities were experimentally validated. Furthermore, conditional gene knockouts were created to investigate the biological importance of the FOH/GGOH salvage pathway. The knockout parasites were viable but more susceptible to fosmidomycin, and their sensitivity to MEP inhibitors could not be rescued by the addition of prenols. Moreover, the knockout parasites lost their ability to use prenols for protein prenylation. These results demonstrate that FOH/GGOH salvage is an additional source of isoprenoids by malaria parasites when *de novo* biosynthesis is inhibited. This study also identifies a novel kind of enzyme whose inhibition may potentiate the antimalarial efficacy of drugs that affect isoprenoid metabolism.

## 1. INTRODUCTION

*Plasmodium falciparum* causes the most severe form of human malaria, a parasitic disease with a high global burden. In 2021, the World Health Organization reported an estimated 247 million cases of malaria and 619,000 malaria-related deaths, with the majority occurring among children and pregnant women in Sub-Saharan Africa. In 2021, 96% of all malaria-related deaths occurred in this region. Resistance to current antimalarial drugs is a significant challenge for malaria control, leading to increased morbidity and mortality (WHO, 2022). Therefore, the identification and development of novel antimalarial therapies are urgently needed.

The most promising targets for the development of antimalarial drugs are those that are unique to the pathogen and not found in humans. The ancestor of apicomplexan parasites underwent endosymbiosis with an alga and thus, possess a non-photosynthetic plastid called the apicoplast (Kohler *et al*, 1997) which contains the targets of some of the current antimalarial drugs in use (Janouskovec *et al*, 2010; Seeber & Soldati-Favre, 2010). The most extensively studied biological process in the apicoplast is isoprenoid biosynthesis via the methylerythritol phosphate (MEP) pathway (Figure 1). Unlike animals, which use the mevalonate (MVA) pathway, the MEP pathway condenses pyruvate and glyceraldehyde 3-phosphate to produce 5-carbon isoprene units, isopentenyl pyrophosphate (IPP) and dimethylallyl pyrophosphate (DMAPP). IPP and DMAPP are enzymatically condensed in geranyl pyrophosphate (10 carbon), farnesyl pyrophosphate (FPP, 15 carbon), and geranylgeranyl pyrophosphate (GGPP, 20 carbon) (Cassera *et al*, 2004; Jordão *et al*, 2011). These metabolites are essential for protein farnesylation and geranylgeranylation (Figure 1). The parasite also produces longer polyprenyl pyrophosphates for the biosynthesis of ubiquinone-8,9 and dolichols, which are mitochondrial cofactors and lipid carriers for sugar transport in protein glycosylation, respectively (Gowda & Davidson, 1999; de Macedo *et al*, 2002; Verdaguer *et al*, 2019; Zimbres *et al*, 2020; Verdaguer *et al*, 2021; Fenollar *et al*, 2022; Okada *et al*, 2022). Isoprenoids produced by the MEP pathway are thus involved in various essential parasitic processes, such as mitochondrial activity and post-translational modification of proteins. The antimalarial drug fosmidomycin inhibits the MEP pathway by targeting the enzyme 1-deoxy-D-xylulose 5-phosphate reductoisomerase (DXR), which converts 1-deoxy-D-xylulose 5-phosphate (DXP) to MEP (Figure 1) (Lell *et al*, 2003). Similarly, classic bacterial ribosome inhibitors (hereafter referred as ribosome inhibitors), such as azithromycin, doxycycline, or clindamycin, can indirectly inhibit isoprenoid biosynthesis by interfering with apicoplast biogenesis. Treated parasites transmit defective organelles to their progeny, leading to a delayed death effect in which the parasites exposed to these drugs cease growth approximately 48 hours after the onset of treatment (Yeh & DeRisi, 2011; Wu *et al*, 2015; Kennedy *et al*, 2019). Until recently, it was assumed that these drugs completely inhibited apicoplast formation and all the metabolic processes associated with this organelle. However, recent studies have found that treatment with ribosome inhibitors leads to the fragmentation of the apicoplast into vesicles. Whereas these vesicles do not maintain active the MEP pathway, they still contain other important enzymes (Swift *et al*, 2021).

**Figure 1.**
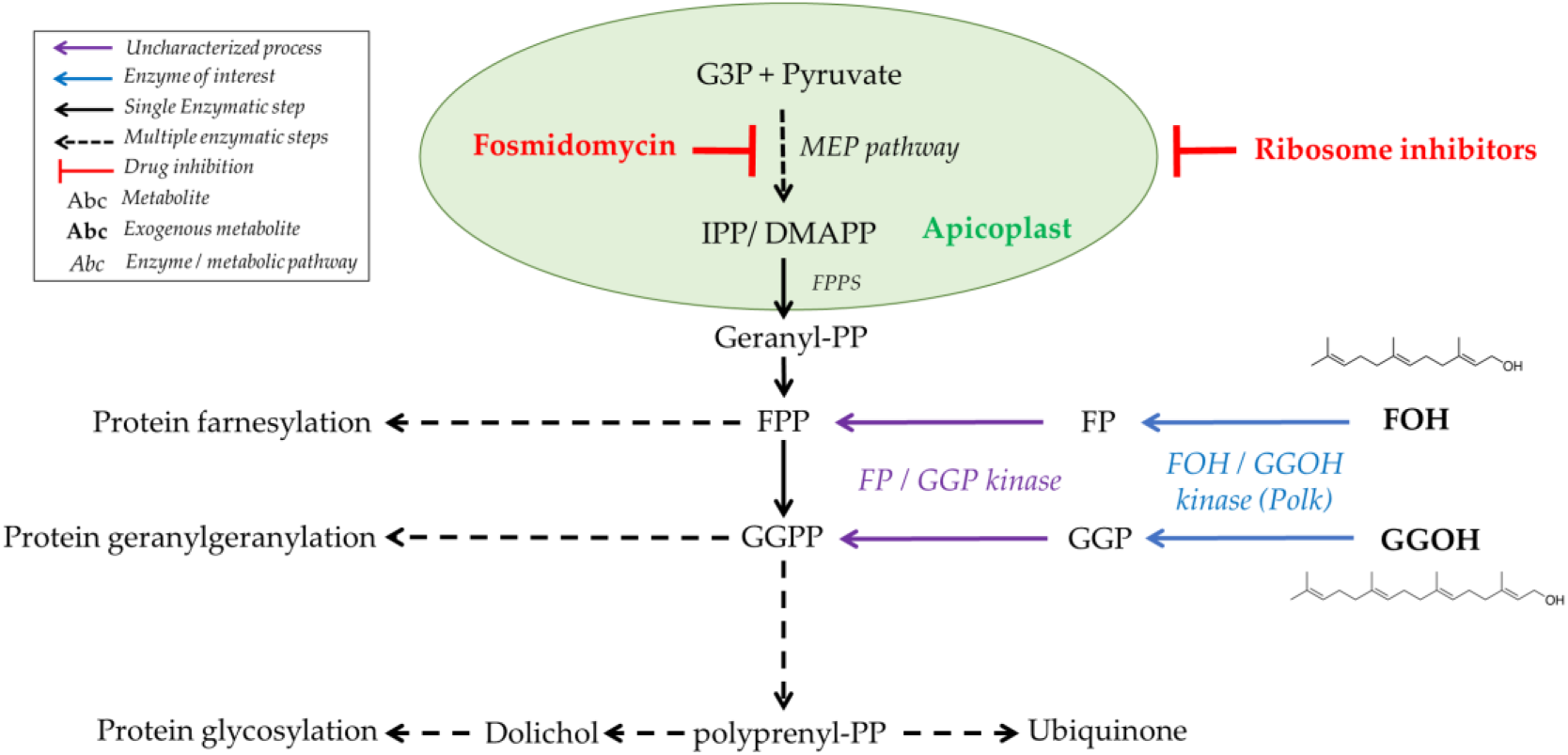
Isoprenoid sources and distribution in malaria parasites. The figure illustrates the sources and distribution of isoprenoids in malaria parasites. The figure includes the biosynthesis of isoprenoids via the MEP pathway, starting with the condensation of glyceraldehyde-3-phosphate (G3P) and pyruvate, and leads to the formation of isopentenyl pyrophosphate (IPP) and dimethylallyl pyrophosphate (DMAPP), and their subsequent condensation to form geranyl pyrophosphate (GPP), farnesyl pyrophosphate (FPP), and geranylgeranyl pyrophosphate (GGPP). These longer isoprenoids are essential for the biosynthesis of ubiquinone, dolichol, and for protein prenylation. The figure also shows the targets of fosmidomycin and ribosome inhibitors in the parasite. The chemical structures of FOH and GGOH are represented.

Parasites can be grown indefinitely *in vitro* when exposed to fosmidomycin or ribosome inhibitors if there is an exogenous source of IPP. Additionally, parasites with impaired isoprenoid biosynthesis can be partially rescued by the addition of FPP/farnesol (FOH) and GGPP/geranylgeraniol (GGOH), but not isopentenol, geraniol, octaprenol, nonaprenol or dolichols (Yeh & DeRisi, 2011; Wu *et al*, 2015; Kennedy *et al*, 2019; Verdaguer *et al*, 2022a). Additional studies characterized the molecular and morphological phenotype of parasites exposed to ribosome inhibitors in order to investigate their isoprenoid requirements. Additionally, metabolomic profiling revealed that the lack of ubiquinone and dolichol biosynthesis is not the primary cause of death, but rather the disruption of the digestive vacuole function (Kennedy *et al*, 2019). It is now understood that the loss of protein prenylation interferes with vesicular trafficking and ultimately affects *P. falciparum*’s feeding, leading to their death. In fact, the sort of prenylated proteins in malaria parasites includes Ras, Rho, and Rap small GTPases, which are involved in cellular signalling and intracellular trafficking (Suazo *et al*, 2016; Verdaguer *et al*, 2022a). Mechanistically, these processes rely on transferases that attach FPP or GGPP moieties to the C-terminal cysteine residues of proteins containing a conserved motif for prenylation, CAAX (C = cysteine, A = aliphatic amino acid, X = diverse terminal residue) (Farnsworth *et al*, 1990).

Several clinical trials using fosmidomycin to treat malaria failed, mostly due to poor antimalarial efficacy (Fernandes *et al*, 2015). Additionally, the combination of fosmidomycin plus clindamycin was also unsuccessful in clinics, with no clear evidence of mutations related to its resistance (Lanaspa *et al*, 2012; Mombo-Ngoma *et al*, 2018). Thus, the failure of these therapies may be due to the intrinsic mechanisms of the parasite or the pharmacokinetics of fosmidomycin (Guggisberg *et al*, 2016). Therefore, it is crucial to conduct further research on isoprenoid metabolism in the parasite to develop effective antimalarial treatments targeting these pathways.

An unresolved matter regarding isoprenoid metabolism in *Plasmodium* is related to classic drug-rescue assays that employ the prenols FOH and GGOH (Zhang *et al*, 2011; Yeh & DeRisi, 2011, Howe *et al*, 2013; Guggisberg *et al* 2014; Kennedy *et al*, 2019; Verdaguer *et al*, 2022a). There is no evidence that prenyl synthases/transferases would preferably bind prenols, or any other polyprenyl derivative, rather than polyprenyl-PP. Prenols may orient parallel to the membrane, while polyprenyl pyrophosphates favour a perpendicular orientation, making their pyrophosphate moieties physically available for interaction with polyprenyl transferases and synthases (Hartley *et al*, 2013; Nakatani *et al*, 2014; Verdaguer *et al*, 2022b). Therefore, a FOH/GGOH salvage pathway was proposed to exist, acting via phosphorylation to incorporate these molecules into the major isoprenoid metabolism (Verdaguer *et al*, 2022b). Our group has previously characterized the transport of FOH and GGOH in *P. falciparum*, and observed that these prenols are phosphorylated, condensed into longer isoprenoids, and incorporated into proteins and dolichyl phosphates (Verdaguer *et al*, 2022a).

Prenol phosphorylation has been biochemically demonstrated to occur in membrane extracts of animal and plant tissues, as well as in archaea (Verdaguer *et al*, 2022b). In 1998, Bentinger *et al*. reported the first evidence of this pathway in mammals, observing the conversion of FOH to farnesyl monophosphate (FP) in the 10,000 x *g* supernatant of rat liver homogenates (Bentinger *et al*, 1998). This FOH kinase activity was located in rough and smooth microsomes and associated with the inner, luminal surface of the vesicles. Further analysis identified an activity capable of phosphorylating FP to FPP. Although the biological function of this pathway remains poorly understood, it is likely to be a mechanism for regulating or bypassing the isoprenoid biosynthetic pathway, recycling isoprenoids released from prenylated metabolite degradation, or even facilitating the use of exogenous isoprenoids (Ischebeck *et al*, 2006; Valentin *et al* 2006; Fitzpatrick *et al*, 2011; Vom Dorp *et al*, 2015; Verdaguer *et al*, 2022b). Biochemical evidence suggest that the pathway is carried out by two separate enzymes: a CTP-dependant prenol kinase (PolK) with FOH/GGOH kinase activities that produce FP or GGP and a polyprenyl-phosphate kinase. However, only a few genes encoding these enzymes have been experimentally identified in plants, including the FOLK gene which encodes a FOH kinase in *Arabidopsis thaliana* (Fitzpatrick *et al*, 2011), the VTE5 gene which encodes a kinase of phytol (a hydrogenated product of GGOH typical from plants) (Valentin *et al*, 2006), and the gene VTE6 which encodes a phytyl-P kinase (Vom Dorp *et al*, 2015). The enzymes responsible for the salvage pathway of prenols in animals and other organisms remain unidentified. The interest in their identification is growing, as recent studies show that prenols can be enzymatically produced by mammal phosphatases (Bansal & Vaidya, 1994; Elsabrouty et al, 2021) or metabolized from dietary sources, playing a role in several diseases (de Wolf *et al*, 2017; Jawad *et al*, 2022). This highlights the potential importance of the FOH/GGOH salvage pathway in limiting the efficacy of treatments which target isoprenoid metabolism.

To address these issues, our efforts focused on identifying the enzymes responsible for the FOH/GGOH salvage pathway in *P. falciparum* as a strategy to improve the efficacy of MEP inhibitors. We identified a gene in malaria parasites that encodes a PolK, experimentally validated for its FOH/GGOH kinase activities. Additionally, using bioinformatics approaches and the creation of conditional gene expression knockout parasites, we sought to understand the biological significance of the FOH/GGOH salvage pathway in malaria parasites. Our findings provide important insights into the isoprenoid metabolism of malaria parasites, and the potential of targeting the FOH/GGOH salvage pathway to improve the efficacy of antimalarial drugs.

## 2. RESULTS

### 2.1 Candidates for apicomplexan prenol kinases are homologous to their plant/algae counterparts and belong to a diverse family with multiple gene duplications

We recently demonstrated the ability of parasites to phosphorylate FOH and GGOH (Verdaguer *et al* 2022a). Hence, we started a bioinformatic search for gene candidates to encode a kinase of prenols. As seeds, we used the *A. thaliana* and *Synechocystis* spp. (strain PCC 6803 / Kazusa) VTE5 proteins (phytol kinase) and *A. thaliana* FOLK protein (farnesol kinase) since their enzymatic activity was already described in the literature (Q9LZ76 and P74653 entries in UniProt, respectively) (Valentin *et al*, 2006; Fitzpatrick *et al*, 2011). Both sequences were defined as control representatives and were used as queries to a survey for homologous sequences in the *P. falciparum* genome in the PlasmoDB database (https://plasmodb.org/). Sequence homology searches were performed using the BlastP algorithm against the protein database of all *Plasmodium* species available (Supporting information, Figure S3). As a result, only two possible orthologous proteins were obtained in *Plasmodium yoelli* genome (PY17X_1224100 and PYYM_1223600). The ortholog of these genes in *P. falciparum* 3D7 and NF54 strains were further identified (PF3D7_0710300 and PfNF54_070015200, with 100% identity between then) and used in multiple alignments with the control representative sequences and other four sequences with putative annotation for phytol kinase (Supporting information, Figure S2). PfNF54_070015200 has 11%, 15% and 10% similarity with *A. thaliana* VTE5, *Synechocystis* spp. VTE5, and *A. thaliana* FOLK protein, respectively. Phylogenetic analysis of the retrieved representative prenol binding proteins (Figure 2A and supporting information Figure S3) display several diverse clades with multiple gene duplications. We discuss five groups, highlighted by coloured boxes in Figure 2A, as follows: a root-external group containing dolichol kinase (DolK) enzymes which are members of the Polyprenol kinase family (InterproScan num. IPR032974; orange, see also Supporting information, Figure S4); phytol/FOH kinases from plants and unicellular algae (PhyK and Folk, respectively, in green, yellow, and cyan); a *Chlamydomonas reinhardtii* clade (A0XX_CHLRE, where A0XX is a generic label for all the sequences from *C. reinhardtii*’s taxa representing different genes) from which specific Apicomplexa monophyletic clade is derived (in red, Supporting information Figure S4). This supports the idea that *Plasmodium*’s PolK is more similar to unicellular algae proteins, which is consistent with the endosymbiosis event that occurred in apicomplexan ancestors.

**Figure 2.**
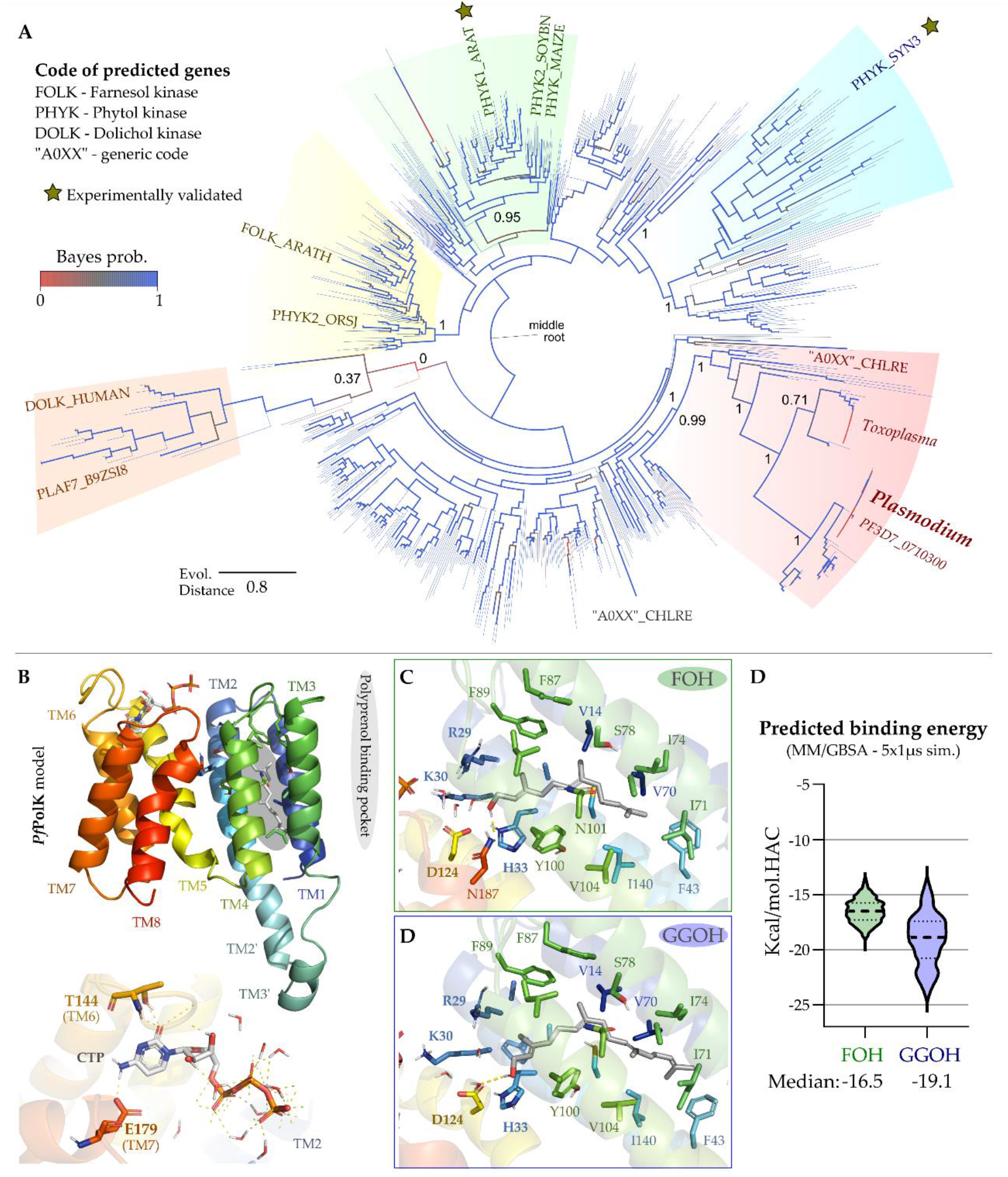
*Pf*PolK phylogenetic analysis and structural model. A) Overall phylogenetic dendrogram of prenol binding proteins generated using maximum likelihood method (see Methods). Branch support values (Bayes posterior probability) are displayed as numbers for the most relevant clade separation, as well as colours (from the highest scores, in blue, to the lowest values, in red) and thickness of the branches. The five discussed groups are highlighted by coloured boxes as follows: root-external group of DolK (orange), phytol/FOH kinases from plants and unicellular algae (PhyK and Folk, respectively, in green, yellow, and cyan) and the *C. reinhardtii* (A0XX_CHLRE, where A0XX is a generic label for all the *C. reinhardtii*’s taxa) clade from which a specific Apicomplexa monophyletic clade arises (in red). Supporting information provides the full phylogenetic tree with all values for branch support and labelled taxa, as well as a key-taxa conversion table. B) AlphaFold 2 *Pf*PolK model displays eight conserved transmembrane helices (TM1-8, coloured) and a potential prenol binding pocket, depicting in the bottom the nucleotide-binding site with the Thre144 and Glu179 composing a hinge region. This model was used to generate the potential binding mode for FOH (green, C) and GGOH (blue, D) by a combination of flexible docking and long molecular dynamics simulations (5×1 µs for each system in explicit solvent and membrane). E) the MD trajectories were utilized to infer the substrates predicted binding energy (Kcal/mol.HAC, where HAC – heavy atom count), suggesting from the median values of the violin-plot displayed distribution that GGOH would have a lower potential binding energy. Dotted lines describe the first quartile amplitude.

We assessed the potential of an AlphaFold-derived structural model of the putative *Pf*PolK to bind prenols using a combination of docking and long Molecular Dynamics (MD) simulations. The generated model displays eight conserved transmembrane helices (TM1-8, Figure 2B), a conserved CTP binding pocket and a potential prenol binding pocket (in grey). The CTP binding pocket ends in the charged clamp motif (Arg29, Lys30 and His33), whose positive charges could be used to orient substrate phosphates into the catalytic conformation, while the nucleotide ring is stabilized by a conserved “hinge” region composed by the main chain of Thr144 and the side chain of Glu179 (Figure 2B, down inset). Meanwhile, the potential prenol binding pocket, composed by the TM’s 1 – 4, can accommodate both FOH (Figure 2C) and GGOH (Figure 2D) relying on the conformational change, upon simulation, of the Phe43 and Ile71 to fit the later. The hydroxyl group of both substrates coordinates the Arg-Lys-His triad by conserved water interactions. The MD trajectories were further utilized to infer the substrates predicted binding energy (Figure 2E), suggesting that GGOH (−19.1 kcal/mol) would have a lower potential binding energy, when compared to the FOH (−16.5 kcal/mol), with both substrates being able to bind *Pf*PolK.

### 2.2 Farnesol/geranylgeraniol kinase activity of *Pf*PolK

To study the catalytic activity of *Pf*PolK candidate, we expressed its gene heterologously in yeast. This expression system was chosen due to its advantages as a eukaryotic protein-expression system, and because it was previously demonstrated that this organism did not phosphorylate FOH and GGOH (Fitzpatrick *et al*, 2011). Therefore, the W303-1A strain of *S. cerevisiae* was transformed with p416-GPD vector (empty vector employed, as a control) or engineered to express *Pf*PolK from p416-P*f*PolK plasmid. Transformant yeasts were grown in SD-an uracil drop-out medium and employed for enzymatic assays. The incubation of yeast extracts with [^3^H] FOH or [^3^H] GGOH plus CTP produced radiolabeled compounds chromatographically compatible with their respective phosphates (Rf ∼0.5). The formation of polyprenyl phosphates was not observed in assays without the addition of CTP or employing wild-type yeasts transformed with the empty vector (Figure 3). No compounds displaying chromatographic compatibility with FPP and GGPP were detected. Although we used the same amounts of substrates in all assays, the chromatographic spots compatible with FOH and FP were more visible than those compatible with GGOH and GGP. This issue is likely caused by varying extraction efficiencies between different prenols.

**Figure 3.**
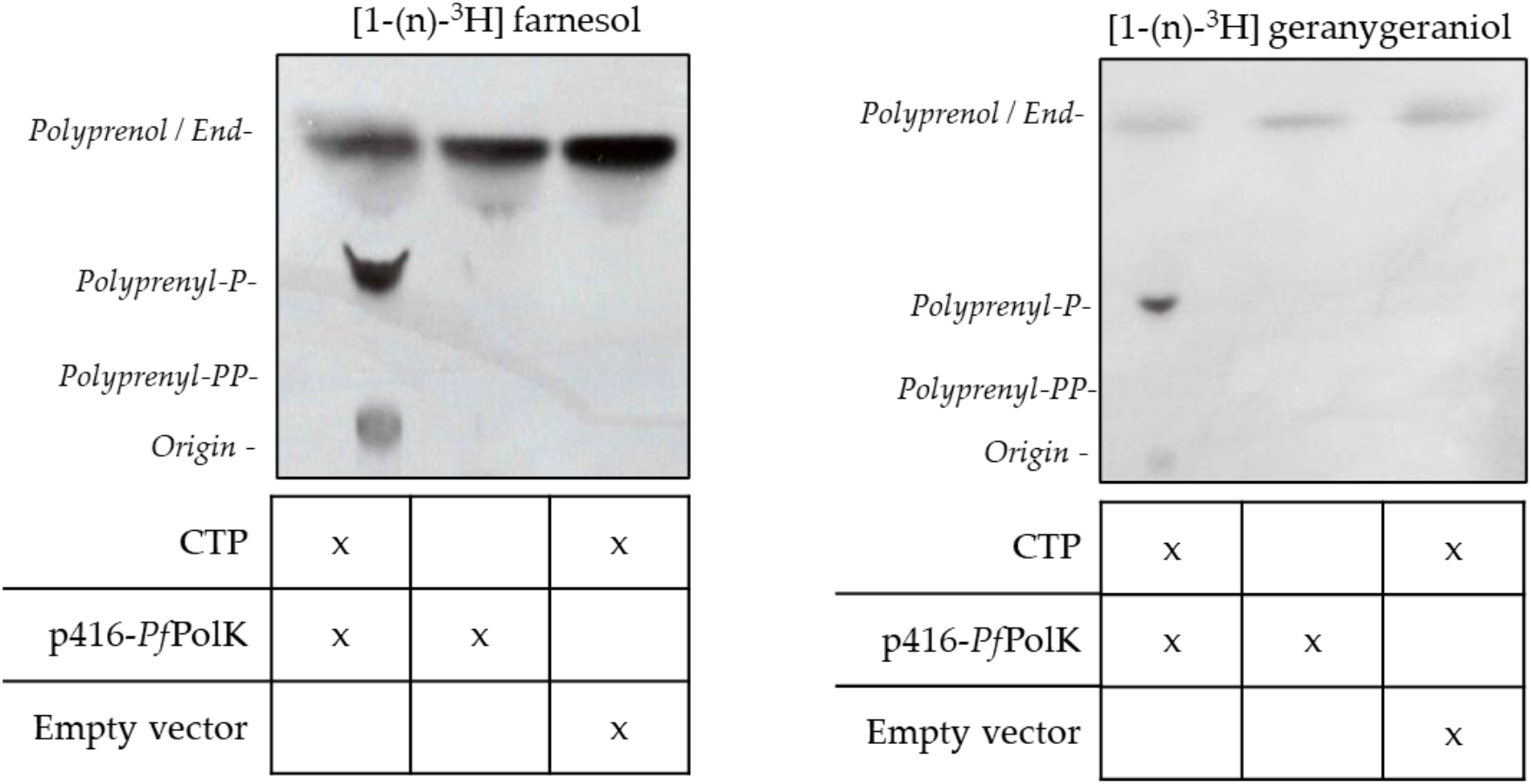
Farnesol and geranylgeraniol kinase activities. Autoradiographies of the PolK enzymatic activity assays using [^3^H] FOH or [^3^H] GGOH as substrates and chromatographed by TLC. The enzyme source of these assays came from whole extracts of yeast strains transformed with either the empty vector (p416-GPD) or p416-*Pf*PolK. Compounds added to the enzymatic reaction are indicated under the TLC autoradiography image. The retention of different standards is also indicated. These experiments were repeated three times with similar results.

### 2.3 Conditional knockout uncovers the relationship between P*f*PolK and MEP inhibitors

After confirming the catalytic activity of *Pf*PolK, we investigated its biological importance in malaria parasites by generating conditional knockout NF54 parasites for PfNF54_070015200 (Figures 4A and 4B). Clones PolK-loxP-C3, E3, and G9 were obtained and genomic integration was confirmed by PCR (Figure 4C). Expected excision of the floxed *Pf*PolK-loxP was also confirmed by PCR 24, 48, 96 and 144 h after knockout induction with rapamycin, generating Δ-PolK parasite lines (Figure 4D). Importantly, Δ-PolK parasites showed no reduction in growth and could be maintained indefinitely in culture, as demonstrated by monitoring their growth for up to 144 hours (Figure 4E). Western blots revealed that transfected parasites had an HA epitope fused to a ∼31 kDa protein, matching the expected size of the PolK protein. In contrast, HA-tagged PolK protein was not detectable in Δ-PolK parasites 48 hours after induction of *Pf*PolK-loxp excision with rapamycin (Figure 4F; see original images in supporting information, Figure S5). Immunofluorescence assays also revealed that the HA-tagged protein did not exclusively co-localize with the apicoplast (Figure 4G), suggesting that the prenol salvage pathway could be at least partially, apicoplast-independent.

**Figure 4.**
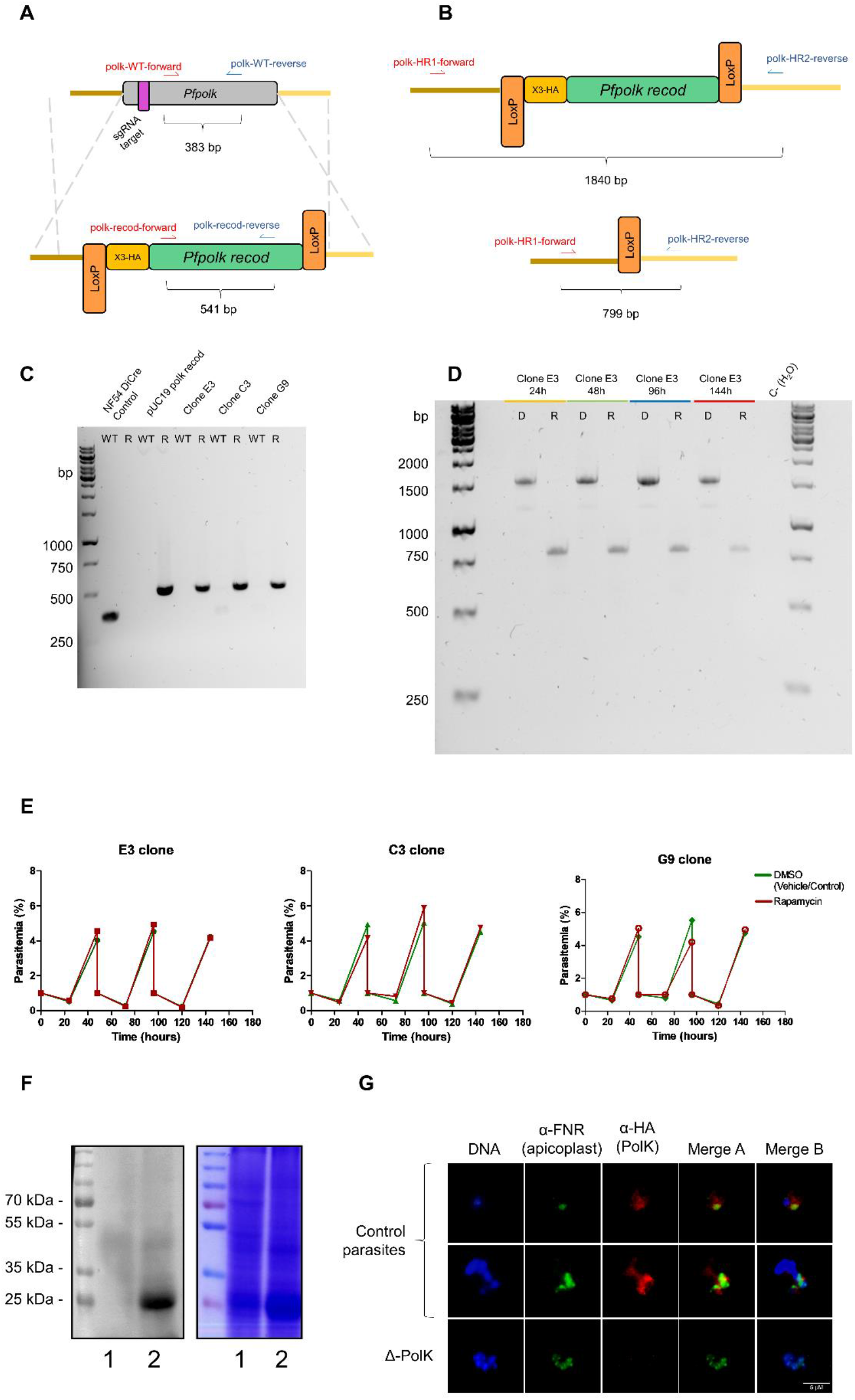
Conditional knockout of *P. falciparum* PolK gene. (A) Diagram depicting the edition of single-exon gene *P. falciparum* PolK. Using Cas9-assisted genome editing, all 612 bp of the native *Pf*PolK (WT) open reading frame were replaced by a recodonized sequence (*Pf*PolK recod) and a 3x-HA sequence (yellow box) in the 5’ end of the sequence and all flanked by two loxP sites (orange boxes). The position of the 20-nucleotide region targeted by the single guide RNA (sgRNA) is indicated (purple box). (B) Diagram of the rapamycin-induced site-specific excision. Recombination between the loxP sites removes the entire recodonized gene (green box) and 3-HA sequence; (C) PCR assessment of the genome integration of the construct in the *Pf*PolK locus. Wild type (NF54 DiCre control) was confirmed using Polk-WT-forward and reverse primers. Using PolK-recod-forward and reverse primers it was detected redoconized version of PolK in transgenic parasites (Clones E3, C3 and G9) and in a plasmid containing the recodonized PolK (pUC19 polk recod). (D) Confirmation of the rapamycin-induced *Pf*PolK gene excision in the clone PolK-loxP E3. The deletion was confirmed by PCR 24, 48, 96 and 144 h after treatment with DMSO (D) or rapamycin (R) using primers polk-HR1-forward and polk-HR2-reverse (in red and blue on panel B, respectively). Excision reduces the amplicon from 1840 bp to 799 bp, disrupting *Pf*PolK. (E) Figure shows the 144h evolution of parasitemia in different clones in which *Pf*PolK gene was excised (parasites exposed to rapamycin) or not (parasites exposed to DMSO). All the graphs represent the mean and SD of at least three experiments. (F) Western blot of transgenic parasites (left) and the respective Comassie stained gel (right). Western blot was performed to analyse the HA-tagged *Pf*PolK of parasites in which *Pf*PolK was excised (lane 1, parasites exposed to rapamycin) or preserved (lane 2, parasites exposed to DMSO). (G) Immunofluorescence analysis of HA-tagged *Pf*PolK of parasites in which *Pf*PolK was excised (parasites exposed to rapamycin, Δ-Polk) or not (control parasites, exposed to DMSO). HA-tagged *Pf*PolK is marked in red, the apicoplast is green and the nucleus in blue.

The susceptibility of these parasites to MEP inhibitors was studied next. Loss of the PolK gene increased the sensitivity of mutant parasites to fosmidomycin two-fold, when compared to wild type parasites (IC_50_ fosmidomycin 1.09 ± 0.33 µM vs 0.51 ± 0.07 µM) (Figure 5A, B). No significant differences were observed in the effect of clindamycin under the same conditions, possibly due wider effects on parasites metabolism. As expected, the presence of prenols, FOH or GGOH, in the medium of control parasites reduced their sensitivity to fosmidomycin. This was evidenced by a 3-fold increase in the IC_50_ for fosmidomycin in the presence of FOH, and a 15-fold increase in the presence of GGOH, when compared to controls growth without prenol supplementation. Likewise, the IC_50_ for clindamycin increased 5-fold in the presence of GGOH, as compared to the control group with no-additions (Figure 5C, D). Remarkably, the addition of prenols did not have any rescue effects on the antimalarial activity of fosmidomycin or clindamycin in Δ-PolK parasites (see Figure 5A-D). This strongly suggests that the presence of PolK is necessary for the rescue effect of prenols on the antimalarial activity of MEP inhibitors.

**Figure 5.**
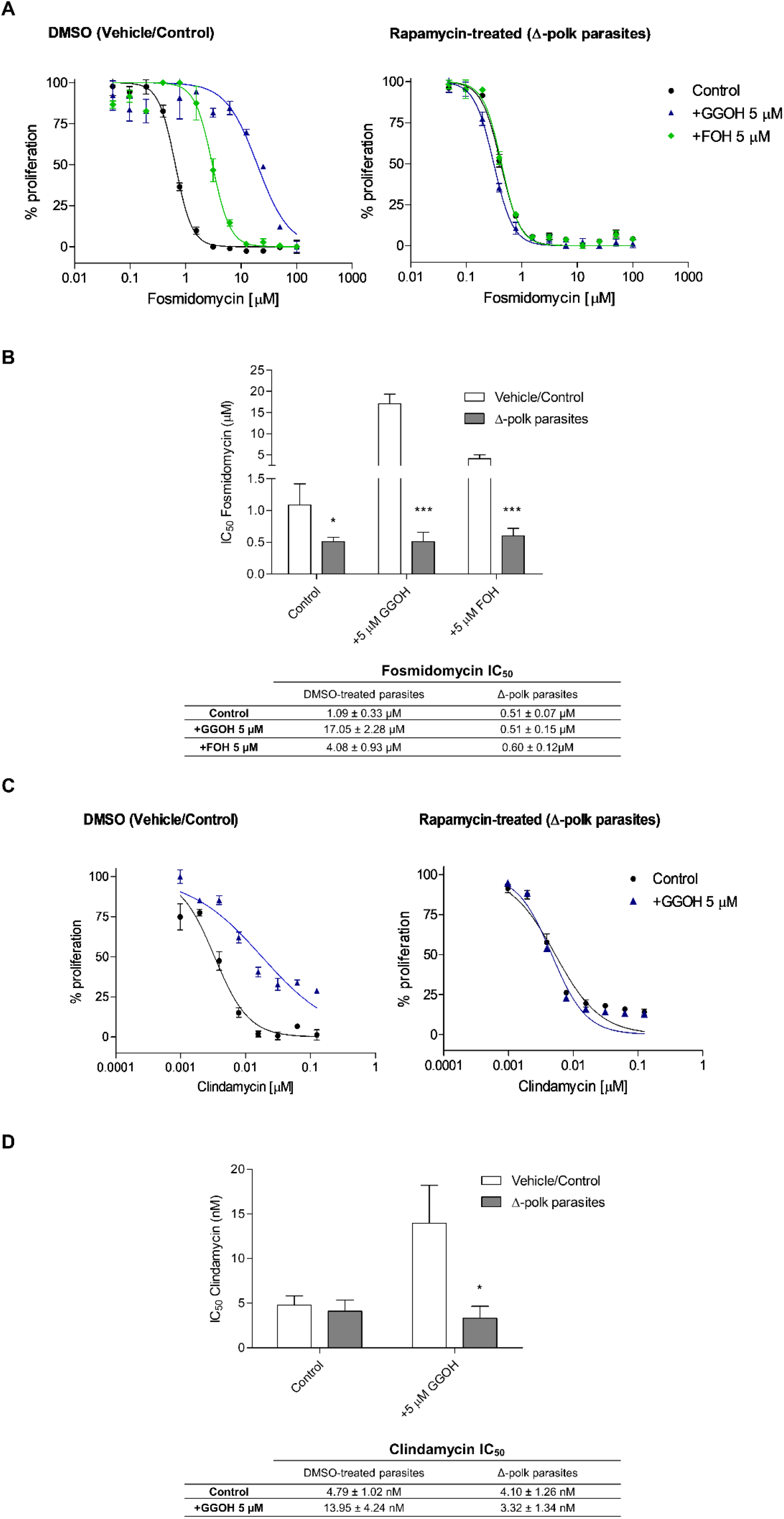
Phenotypic characterization of knockout parasites. (A) Fosmidomycin dose-response curves after 48h of parasites maintaining a functional *Pf*PolK (DMSO (Vehicle/control)) or Δ-PolK parasites. These parasites were cultured in RPMI medium in the presence or absence of the indicated prenols (5 µM). (B) Fosmidomycin IC_50_ values of the results exposed in the previous panel. (C) Clindamycin dose-response curves after 96 h parasites maintaining a functional *Pf*PolK (DMSO (Vehicle/control)) or Δ-PolK parasites. These parasites were cultured in RPMI medium in the presence or absence of GGOH (5 µM), as indicated. (D) Clindamycin IC_50_ values of the results exposed in the previous panel. Statistical analysis was made using one-way ANOVA/Dunnet’s Multiple Comparison Test.*p<0.05, **p<0.01, ***p<0.001. Comparison made to Vehicle/Control data. Error bars represent standard deviation (n = 3).

### 2.4 Farnesol and geranylgeraniol phosphorylation is required for their utilization for protein prenylation

As mentioned above, lack of protein prenylation disrupts the function of the digestive vacuole and leads to the loss of parasitic homeostasis (Kennedy et al., 2019). To understand the effects of PolK deletion on FOH utilization, the incorporation of [^3^H] FOH and [^3^H] isoleucine (control) into proteins was determined in parasites with functional *Pf*PolK and after the excision of the gene (Figure 6). The deletion of *Pf*PolK resulted in a significant decrease in counts per minute (CPM) corresponding to [^3^H] FOH-labeled proteins compared to parasites with an intact PolK enzyme. Only a few counts were still detected in Δ-PolK parasites, likely corresponding to the remaining dolichol-P oligosaccharide and/or other radiolabeled lipids that were not covalently bound to proteins. It is worth noting that all parasites incorporated similar levels of [^3^H] isoleucine into proteins, indicating no observable defects in protein synthesis other than prenylation.

**Figure 6.**
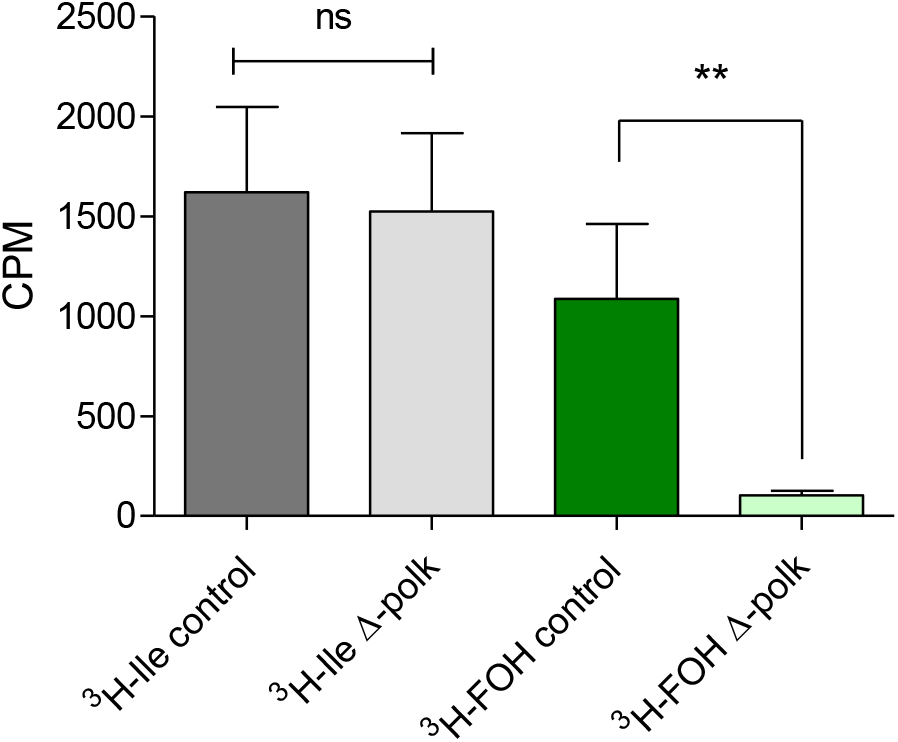
Farnesol and isoleucine incorporation into proteins in Δ-PolK parasites. The graph shows the levels of incorporation of ^3^H-FOH and ^3^H-isoleucine (^3^H-Ile) into TCA-precipitated proteins in parasites maintaining or not a functional *Pf*PolK. Statistical analysis was made using one-way ANOVA One-way ANOVA / Tukey’s Multiple Comparison Test.*p<0.05, **p<0.01, ***p<0.001. Comparison made to between samples of parasites exposed to the same radiolabelled precursor but maintaining or not *Pf*PolK. Error bars represent standard deviation (n = 3).

## 3. DISCUSSION

Malaria parasites can grow indefinitely in the presence of fosmidomycin or ribosome inhibitors if exogenous IPP is added to the culture medium (Yeh & DeRisi, 2011). Moreover, pharmacologically-induced isoprenoid biosynthesis deficiency can be transiently mitigated by the addition of FOH, GGOH and unsaponifiable lipids from food, such as sunflower oil and arugula (Yeh & DeRisi, 2011; Wu *et al*, 2015; Kennedy *et al*, 2019; Verdaguer *et al*, 2022a). In a recent study, our group showed that radiolabelled FOH and GGOH can be incorporated into various long-chain prenols (>20C in length), dolichols, and proteins (Verdaguer *et al*, 2022a). However, since all characterized polyprenyl transferases and synthases use polyprenyl pyrophosphates as their natural substrates, and there is no evidence that these enzymes can use prenols, a phosphorylation pathway for FOH and GGOH salvage is required in malaria parasites. Therefore, we hypothesized the existence of a phosphorylation pathway for FOH and GGOH salvage in malaria parasites and, remarkably, we recently described that *P. falciparum* phosphorylate [^3^H] FOH and [^3^H] GGOH into their pyrophosphate counterparts. These results significantly support the existence of a plasmodial FOH/GGOH salvage pathway (Verdaguer et al 2022a). In photosynthetic organisms the phosphorylation of FOH and GGOH is carried out by two separate enzymes: a prenol kinase and a polyprenyl-phosphate kinase (Verdaguer *et al* 2022a). In photosynthetic organisms the phosphorylation of FOH and GGOH is carried out by two separate enzymes: a prenol kinase and a polyprenyl-phosphate kinase (Verdaguer et al 2022b). However, only a few genes have been unequivocally identified to encode those enzymes (Valentin et al 2006; Fitzpatrick et al, 2011; Vom Dorp et al, 2015). Despite the scarce literature on prenol kinases, we were able to identify a candidate for PolK in *P. falciparum* through BLAST analysis. This enzyme was heterologously expressed in *S. cerevisiae* and its FOH and GGOH kinase activity was biochemically confirmed. Remarkably, *Pf*PolK only catalyses lipid mono-phosphorylations, although it can use either FOH or GGOH as lipid substrates. Consequently, the parasite probably possesses at least another enzyme with polyprenyl-P kinase activity to make prenols available for further use. The low sequence similarity shared by these enzymes is probably the reason for not finding it in the initial searches.

The results here presented could represent a significant contribution to the field of isoprenoid metabolism, as it links the plasmodial prenol kinase function to a new class of enzymes in nature. Remarkably, this is the first FOH/GGOH kinase enzyme discovered in a non-photosynthetic organism. As observed in the phylogenetic analysis, putative prenol kinases comprise a very divergent group of clades, including the group of DolK, phytol/FOH kinases from plants and unicellular algae, and a *C. reinhardtii* clade, from which a specific Apicomplexa monophyletic clade is derived. This finding supports the idea that *Plasmodium*’s PolK is more similar to unicellular algae proteins, consistent with the endosymbiosis event that occurred in apicomplexan ancestors. Also, the paucity of verified enzymes in all commented clades open the possibility to assign pyrophosphorylating activity to some of these putative enzymes. The model provided structural insights into *Pf*PolK, revealing eight conserved transmembrane helices and a conserved CTP binding pocket that could be used to orient the phosphates into the catalytic conformation. Additionally, the nucleotide ring is stabilized by a conserved hinge region. The model also uncovered a novel prenol binding pocket composed of TM’s 1-4, capable to accommodate both FOH and GGOH.

After *Pf*PolK identification, we studied the biological relevance of the FOH/GGOH salvage pathway. *Pf*PolK appears to be expressed throughout all stages of asexual intraerythrocytic development (Chappell *et al*, 2020), highlighting its potential importance in the parasite’s lifecycle. Previous studies indicated that the gene is essential in *P. falciparum* (Zhang *et al*, 2018), whereas studies in rodent malaria parasites predicted it to be dispensable (Bushell *et al*, 2017). Interestingly, Δ-PolK parasites remained fully viable with no observable growth rate changes, in agreement with the observations made in rodent parasites. These results suggest that while the FOH/GGOH salvage pathway may be an important alternative source of isoprenoids, it is not essential for parasite survival, probably due to the endogenous biosynthesis of isoprenoids via the MEP pathway. IFAs revealed that *Pf*PolK did not exclusively co-localizes with the apicoplast marker (Figure 4G), suggesting that the prenol salvage pathway is at least partially, apicoplast-independent. This contradicts previous large-scale studies which target this protein to the apicoplast (Boucher *et al*, 2018). In line with our findings, bioinformatic deep learning signal prediction indicates that this enzyme is localized in the endoplasmic reticulum of malaria parasites (Thumuluri *et al*, 2022). In fact, other studies indicate that the majority of prenol kinase activities in animals and bacteria are located in the approximately 10,000 x *g* supernatant of tissue homogenates, rough and smooth microsomes, and associated with the inner, luminal surface of the vesicles (Verdaguer *et al*, 2022b).

Δ-PolK parasites could not be rescued from the antimalarial effects of fosmidomycin or clindamycin by the addition of FOH or GGOH. As far as we know, this observation is the first evidence of FOH and GGOH phosphorylation as a mandatory step for their utilization by living organisms, highlighting the biological relevance of the FOH/GGOH salvage pathway particularly when *de novo* isoprenoid biosynthesis is inhibited. The biological relevance of the FOH/GGOH salvage pathway in *P. falciparum* raises important questions. In our view, this pathway could serve as an alternative source of isoprenoids for the parasite, independent of the Apicoplast. Furthermore, the pathway could be important for recycling prenols released from the degradation of endogenous prenylated metabolites. This mechanism is similar to the prenol salvage pathway in plants, which also recycles prenols from chlorophyll degradation (Ischebeck *et al*, 2006; Valentin *et al* 2006). In addition, *P. falciparum* FOH/GGOH salvage pathway in the parasite may serves as a mechanism to scavenge FOH and GGOH derived from the host. Thus, prenol scavenging could help the parasite optimize its energy usage for isoprenoid biosynthesis and partially complement its metabolic requirements. Remarkably, the fact that these nutrients can reduce the efficacy of MEP inhibitors *in vitro* suggests the potential importance of this pathway in limiting the effectiveness of this type of antimalarial drug in clinical settings (Verdaguer *et al*, 2022a). Contrarily to this hypothesis, other authors previously concluded that the parasite may rely exclusively on endogenous isoprenoid biosynthesis based on the observation that increasing concentrations of human plasma components in culture did not affect the antimalarial effect of fosmidomycin (Yeh & DeRisi, 2011). However, it is important to note that the concentration of prenols in blood and the physiology of their absorption/excretion in humans remain largely unknown (based on data from the Human Metabolome Database site, http://www.hmdb.ca/). Thus, further research is needed to assess if the FOH/GGOH salvage pathway naturally scavenges host prenols during parasite infections and to determine whether this phenomenon could be related to the limited efficacy of fosmidomycin in clinical trials. The identification of *Pf*PolK opens up new avenues for these studies to be conducted, and may ultimately lead to the development of novel strategies for combatting malaria. Notewothy, Δ-PolK parasites showed to be slightly more susceptible to fosmidomycin. In our point of view, this observation may be a consequence of the presence of small amounts of FOH and/or GGOH in the *in vitro* culture system, may be coming from the bovine components or human red blood cells (note that no data is available about FOH/GGOH quantification in blood or commercial serum substitutes). While this source may not be sufficient to supply the parasite’s isoprenoid requirements *in vitro* under normal culture conditions, it may still contribute to some extent to the observed effects of fosmidomycin *in vitro* tests.

Besides malaria, the dietary consumption of GGOH found in foods like vegetable oils has already been shown to have implications for cancer therapy. Studies reveal that dietary GGOH limits the efficacy of statins, commonly used inhibitors of the mevalonate pathway, in treating certain types of cancers. Specifically, GGOH-rich foods can block statin-induced regression of ovarian tumour xenografts in mice. These findings show that dietary prenols are metabolized and have a significant impact on the outcome of clinical trials for cancer therapies (de Wolf *et al*, 2017; Prior *et al*, 2012; Healy *et al*, 2022). The use of isoprenoid biosynthesis inhibitors has also been explored in other MEP pathway-dependent pathogens such as *Toxoplasma* (Nair *et al*, 2011), *Babesia* (Wang *et al*, 2020), and *Mycobacterium tuberculosis* (Brown & Parish, 2008), as well as in MVA pathway-dependent parasites such as *Leishmania* (Dinesh *et al* 2014). Importantly, most of these pathogens can also be rescued from fosmidomycin by FOH and GGOH (Zhang *et al*, 2011; Li *et al*, 2013; He *et al*, 2018; Kennedy *et al*, 2019; Wang *et al* 2020) and thus, possibly possess an active prenol salvage pathway. To date, isoprenoid biosynthesis inhibitors have only been used in clinical settings for their cholesterol-lowering effects (e.g. simvastatin) and to prevent bone resorption in osteoporosis (e.g. FPP synthase inhibitors such as bisphosphonates) (Verdaguer *et al*, 2022b). However, available data suggest that these inhibitors might have potential in the treatment of several infectious diseases and cancer. The identification and characterization of *Pf*PolK as an enzyme critical for the FOH/GGOH salvage pathway not only provide new insights into the mechanisms underlying malaria parasite metabolism, but also opens new avenues for the utilization and improvement of antimalarial therapies currently under study.

## 4. CONCLUSIONS

The focus of this work was the identification of the enzymes responsible for the salvage pathway of FOH and GGOH in the parasite and their relationship with MEP-targeting drugs. As a result, we identified *Pf*PolK, a novel lipid kinase. Through biochemical and molecular approaches, the catalytic activity and biological importance of this transmembrane enzyme were characterized. Our data revealed the non-essential role of *Pf*PolK in parasite survival and its crucial involvement in the use of exogenous prenols for protein prenylation. *Pf*PolK is also key for maintaining cell homeostasis under the effects of MEP inhibitors. Indeed, we think the findings of this study are not only relevant to understand the fascinating metabolism of malaria parasites, but also to provide new insights into the evolution of the isoprenoid metabolism, and possibly to for the development of novel therapeutic strategies in the treatment of other diseases.

## 5. MATERIALS AND METHODS

### 5.1 Reagents, stock solutions and parasitic strains

AlbuMAX™ II Lipid-Rich BSA and RPMI-1640 were purchased from Thermo Fisher Scientific^®^ (Leicestershire, UK). Dolichol and dolichyl-P 13-21 were purchased from Avanti^®^ (Alabama, USA). [1-(n)-^3^H] GGOH (14 Ci/mmol; 1 mCi/mL), [1-(n)-^3^H] FOH (14 Ci/mmol; 1mCi/mL) and L-[4,5-^3^H(N)] isoleucine (30-60 Ci/mmol; 1 mCi/ml) were purchased from American Radiolabeled Chemicals^®^ (St. Louis, USA). SYBR Green I^®^ nucleic acid gel stain and SYTO® 11 were purchased from Thermo Fisher Scientific^®^ (Waltham, Massachusetts, EUA). Sterile stock solutions were prepared at 10 mM for fosmidomycin sodium salt hydrate in water, 2 mM clindamycin hydrochloride in water, 125 mM of GGOH in ethanol and 200 mM of each other non-radiolabelled prenols in ethanol. All other reagents were purchased from Sigma^®^ (St. Louis, Missouri USA) or specific companies, as cited in the text. Polyprenyl phosphates were obtained by mild acid treatment of the respective commercial pyrophosphates (Sigma) (Ohnuma *et al*, 1996). For this work, a Cre-LoxP *P. falciparum* NF54 strain (Tibúrcio et al., 2019), a generous gift of Moritz Treeck (The Francis Crick Institute, London, United Kingdom), was employed.

### 5.2 *P. falciparum in vitro* culture and synchronization

*P. falciparum* NF54 DiCre cells were cultured *in vitro* following the Trager and Jensen culture method employing RPMI-1640 medium completed with 0.5% AlbuMAX™ II Lipid-Rich BSA. Parasites were maintained in 75 cm^2^ cell culture flasks at 37 °C (Trager & Jensen, 1976; Radfar *et al*, 2009; Crispim *et al*, 2022). The culture medium pH was adjusted to 7.4 and was introduced a gas mixture of 5% CO_2_, 5% O_2_ and 90% N_2_ purchased from Air Products Brasil LTDA^®^ (São Paulo, SP, Brazil). Parasite synchronization at ring stage was performed with 5% (w/v) D-sorbitol solution as described previously (Lambros & Vanderberg, 1979). Parasite development was monitored microscopically on Giemsa-stained smears. PCR for mycoplasma and optic microscopy were used to monitor culture contamination (Rowe *et al*, 1998)

### 5.3 Metabolic labelling of parasites

Our work focused on biochemical experiments in schizont stages because of previous studies showing a higher incorporation rate of [^3^H] isoprenic moieties at this stage (Kimura *et al*, 2011). For this, synchronous cultures of *P. falciparum* at the ring stage in 25 cm^2^ flasks were labelled with either 0.75 μCi/ml [^3^H] FOH or 40 μCi/ml [^3^H] isoleucine employed as control of protein synthesis (Martin & Kirk, 2007). After 12–16 h, parasites at trophozoite/schizont stages were obtained by saponin lysis (Christopher & Fulton, 1939). For this, cultures pellets were lysed with 30 mL 0.03 % saponin in PBS at 4 °C. Parasites were then centrifuged at 1,500 *x g* for 5 min at 4 °C and subsequently washed in PBS.

### 5.4 Assessment of radiolabelled proteins

The assessment of radiolabeled proteins was performed following a similar protocol as described elsewhere (Buesing & Gessner, 2003). Radiolabeled parasites were suspended in 100 µL of lysis buffer (2% w/v SDS, 60 mM DTT in 40 mM Tris-Base pH 9). The samples were then cooled at room temperature, and proteins were precipitated by adding 20% trichloroacetic acid (TCA) in acetone at 4 °C. The samples were kept on ice for 5 minutes, and the proteins were collected by centrifugation at 12,000 × g for 10 minutes. The precipitate was washed three times with 80% acetone. Subsequently, the proteins were dissolved by incubating them at 90 °C in alkaline buffer (0.5 M NaOH, 25 mM EDTA, 0.1 w/v SDS in water) for 30 minutes. Finally, 1 mL of liquid scintillation mixture (PerkinElmer Life Sciences, MA, USA) was added to the samples. After vortex, the radioactivity of samples was measured using a Beckman LS 5000 TD β-counter apparatus (Beckman, CA, USA) and results were analysed using GraphPad Prism® software.

### 5.5 Drug-rescue assays in malaria parasites

In some cases, it was calculated the dose–response curve and the concentration of drug/metabolite required to cause a 50% reduction in parasite growth (IC_50_ value). Assays started at the ring stage at 1% or 0.15% parasitemia and had a duration of 48 h or 96h. Serial dilutions of the antimalarials were prepared in 96-well microplates in RPMI complete medium supplemented or not with FOH / GGOH. Solvent controls and untreated controls were always included and results were analysed by GraphPad Prism^®^ software. All experiments which monitor parasitic growth were performed at least three times with three or four technical replicates. Parasitemia was monitored by flow cytometry using the nucleic acid stain SYTO 11 (0.016 μM) (Life Technologies no. S7573) in a BD LSRFortessa machine as previously described (Portugaliza *et al*., 2019) or in a BD FACSCalibur machine as previously described (Rovira-Graells *et al*., 2016). The data was adjusted to a dose–response curve to determine the IC_50_ value.

### 5.6 Bioinformatics

#### 5.6.1 Sequence similarity search and phylogenetic tree

Sequences from model organisms were retrieved from UniProt, using the term ‘prenol kinase’ as the keyword. Sequences were retrieved from NCBI/GenBank using the Blast tool (with scoring matrix BLOSUM45 for distant similar sequences) with an e-value cut-off of 10^-5^ creating a dataset. Additionally, no similar sequences were found in vertebrate genomes. Sequence renaming and editing were performed with in-house Perl scripts. Sequences with less than 30% global similarity or missing the ORF initiation codon were excluded from further analyses. The full dataset was clustered by similarity (70%) using CD-Hit (Huang *et al*, 2010) and a set of representative sequences were selected for global alignment using Muscle (Edgar, 2004). This algorithm often selects single organisms representing a full clade of highly similar sequences, randomly selecting a centroid sequence within the cluster as a representative.

Maximum likelihood phylogenetic tree was generated using PhyML 3.0 (Guindon *et al*, 2010), with posterior probability values (aBayes) as branch statistical support. The substitution model JTT was selected for calculations, by ProtTest3 (Darriba *et al*, 2011), based on the highest Bayesian Information Criterion values. All other parameters, except the equilibrium frequencies, were estimated from the dataset. Dendrogram figures were generated using FigTree 1.4.4 (see http://tree.bio.ed.ac.uk/software/figtree/", last access in April 2023).

#### 5.6.2 Alphafold model and molecular docking

*Pf*PolK model was retrieved from AlphaFold database (sequence: PF3D7_0710300, https://alphafold.ebi.ac.uk/entry/C0H4M5) and prepared using the PrepWizard implemented in Maestro 2022v4 with standard options. All substrate ligands for docking were drawn using Maestro and prepared using LigPrep to generate the three-dimensional conformation, adjust the protonation state to physiological pH (7.4), and calculate the partial atomic charges, with the force field OPLS4. Docking studies with the prepared ligands were performed using Glide (Glide V7.7), with the flexible modality of Induced-fit docking (Sherman *et al*, 2006; Friesner *et al*, 2006) with extra precision (XP), followed by a side-chain minimization step using Prime. Ligands were docked within a grid around 13 Å from the centroid of the orthosteric pocket, identified using SiteMap (Schrödinger LCC) (Halgren, 2009), generating ten poses per ligand. Docking poses were visually inspected, independently from the docking score, and those with the highest number of consistent interactions were selected for simulation.

#### 5.6.3 Molecular dynamics simulations

*Pf*PolK model with the different substrates was simulated to clarify which residues contributed to the stability within the binding site. Molecular Dynamics (MD) simulations were carried out using the Desmond engine (Bowers *et al*, 2006) with the OPLS4 force-field (Lu *et al*, 2021). The simulated system encompassed the protein-ligand/cofactor complex, a predefined water model (TIP3P) (Jorgensen *et al*, 1983) as a solvent, counterions (Na^+^ or Cl^-^ adjusted to neutralize the overall system charge) and a POPC membrane based in the transmembrane motifs determined in the model. The system was treated in an orthorhombic box with periodic boundary conditions specifying the shape and the size of the box as 10×10×10 Å distance from the box edges to any atom of the protein. Short-range coulombic interactions were performed using time steps of 1 fs and a cut-off value of 9.0 Å, whereas long-range coulombic interactions were handled using the Smooth Particle Mesh Ewald (PME) method (Darden *et al*, 1993). PolK+GTP+substrates systems were then subjected to simulations of 100 ns for equilibration purposes, from which the last frame was used to generate new replicas. The equilibrated system underwent at least 1 µs production simulation, in four-five replicas (total of 5 µs per substrate), followed by analysis to characterize the protein-ligand interaction. The results of the simulations, in the form of trajectory and interaction data, are available on the Zenodo repository (code: 10.5281/zenodo.7540985). MD trajectories were visualized, and figures were produced using PyMOL v.2.5.2 (Schrödinger LCC, New York, NY, USA).

Protein-ligand interactions and distances were determined using the Simulation Event Analysis pipeline implemented using the software Maestro 2022v.4 (Schrödinger LCC). The compounds’ binding energy was calculated using the Born and surface area continuum solvation (MM/GBSA) model, using Prime (Jacobson *et al*, 2004) and the implemented thermal MM/GBSA script. For the calculations, each 10^th^ frame of MD was used. Finally, root mean square deviation (RMSD) values of the protein backbone were used to monitor simulation equilibration and protein changes (supporting information Figure S1). The fluctuation (RMSF) by residues was calculated using the initial MD frame as a reference and compared between ligand-bound and apostructure simulations (supporting information Figure S2).

### 5.7 Generation of conditional Δ-*polk* parasites

A single guide RNA (sgRNA) targeting the PolK genomic locus in PfNF54 strain (PfNF54_070015200) was designed with the CHOPCHOP gRNA Design Tool (Labun *et al*, 2019). To generate the plasmid expressing the *Streptococcus pyogenes* Cas9 and the sgRNA, the primers 5′-AAGTATATAATATTGGACATAGAACAATGTCACAAGTTTTAGAGCTAGAA-3′ and 5′-TTCTAGCTCTAAAACTTGTGACATTGTTCTATGTCCAATATTATATACTT-3′ were annealed and ligated into a BbsI-digested pDC2-Cas9-hDHFRyFCU plasmid (a gift from Ellen Knuepfer) (Knuepfer *et al*, 2017). The donor plasmid, a pUC19 plasmid containing the recodonized PfNF54_070015200 gene, was manufactured by GenWiz Gene Synthesis (Azenta, Chelmsford, USA) with the coding sequence for a 3xHA (Human influenza hemagglutinin) tag in the N-terminal part of the PolK. In addition, this sequence was flanked with two loxP sequences and two homology regions of 500 bp corresponding to intergenic regions upstream and downstream of PfNF54_070015200. Confirmation of the appropriate modification of the PolK gene after transfection was assessed by diagnostic PCR using primers that specifically recognized the wild-type sequence (PolK-WT-forward: 5’-GGATATAGGAGAGGTTTGCCAC-3’ and PolK-WT-reverse: 5’-CCTACTATTGCCGCCATTG-3’) or the recodonized sequence (PolK-recod-forward: 5’-GCTTCGTATTGTTCGTGATA-3’ and PolK-recod-reverse: 5’-CCACCGAACAACTCTAAGAA-3’). Subsequent limiting dilution was performed to generate clones of PolK locus-modified parasites resulting in the isolation of PolK-loxP-C3, PolK-loxP-E3 and PolK-loxP-G9 clones, in which the modification of the locus was reconfirmed again by PCR. Efficiency of the conditional excision of the floxed PolK-loxP clones was assessed by the addition of 50 nM rapamycin or dimethyl sulfoxide (DMSO; vehicle control) in ring-stage synchronized cultures. Cells were treated for 24 h, followed by washing and incubation for another 24 h to allow parasite maturation. To demonstrate the efficient excision of PolK-loxP, gDNA from the clones were obtained and used in diagnostic PCR with primers annealing in the homology regions PolK-HR1-forward (5’-ATGATATTTACCATAATTTATGGGC-3’) and PolK-HR2-reverse (5’-CTGTTTTTTCTCTTTATTTCCTTCTC-3’). The three clones were used in all subsequent experiments.

### 5.8 Immunofluorescence assays

Before the immunofluorescence assays (IFA) procedure, μ-Slide eight-well chamber slides (Ibidi GmbH, Gräfelfing, Germany) were incubated in a working poly-L-lysine solution (1:10 dilution from stock 0.1%) for 5 minutes at RT. The poly-L-lysine was then removed with suction and the slides were left to dry. In parallel, parasite cultures were washed 3 times with RPMI medium and 150 μL of culture were placed in the pre-treated slide. The cultures were then fixed by adding 150 μL of paraformaldehyde 4% in PBS to the slides and incubating them at 37 °C for 30 minutes. After washing the cultures once with PBS, the cultures were permeabilized by adding 150 μL of 0.1% Triton-X-100 in PBS and incubating them at room temperature for 15 minutes. The cultures were then washed 3 times with PBS, and were blocked by adding 150 μL of 3% BSA in PBS and incubating them for 30 minutes at room temperature at 400 rpm, orbital agitation. The cultures were then washed 3 times with PBS and 150 μL of primary antibody solution (Rabbit polyclonal anti-ferredoxin-NADP reductase, diluted 1:100 and Rat Anti-HA, diluted 1:20 in 0.75% BSA/PBS) was then added, and the cultures were incubated overnight at 4°C with at 400 rpm. Afterwards, the cells were washed 3 times with PBS to remove the excess primary antibody solution. The supernatant was removed and secondary antibody solution (Goat anti-Rabbit IgG (H+L) Alexa Fluor 488, #A11034, (Life Technologies, Carlsbad, California, EUA) and Goat anti-Rat IgG (H&L) - AlexaFluor™ 594, #A11007 (Invitrogen, Waltham, Massachusetts, EUA), diluted 1:100 in 0.75% BSA/PBS, was added to the cultures and incubated for 1 hour at room temperature at 400 rpm, orbital agitation. Hoechst (Thermo Fisher Scientific, Waltham, Massachusetts, EUA) was diluted 1:1000 in the mix and also added to cultures. The cells were then washed 3 times with PBS to remove excess secondary antibody. The supernatant was removed and the slides maintained in 150 μL of PBS. For microscopy analysis, an Olympus IX51 inverted system microscope, equipped with an IX2-SFR X-Y stage, a U-TVIX-2 camera, and a fluorescence mirror unit cassette for UV/blue/green excitation and detection was employed.

### 5.9 Western blot

Cultures with 5% parasitaemia at trophozoite/schizont stages were centrifuged in a 15 mL tubes, resuspended in 2 volumes of 0.2% saponin in PBS and incubated on ice for 10 min. Then, 10 mL of PBS was added to each sample and the mixture was centrifuged at 1800 x *g* for 8 min at 4 °C. The supernatants were removed and the saponin treatment was repeated two more times. The pellets were transferred to a 1.5 mL vial and washed with PBS, then resuspended in 100 µL of lysis buffer. BioRad Bradford Assay was carried out and 10 µg of each sample was applied in SDS-PAGE gels and then transferred to a PVDF membrane (Bio-Rad, 0.45 µm pore size) by electro-transfer (30 V constant overnight) in the Mini Trans-Blot cell module (Bio-Rad). The membrane was blocked for 1 h at 4 °C with 3% (w/v) BSA in PBS-T (10 mM Tris-HCl pH 8.0, 0.05% Tween 20) and then incubated for 1 h with a rat anti-HA primary antibody (1:500 [vol/vol] in PBS-T) for HA-detection. After three washing steps PBS-T, a secondary goat anti-rat IgG antibody, HRP conjugated, was used at 1:1000 and incubated for 1 hour. Following three washing steps, the membrane is developed with a chemiluminescent substrate (Super SignalTM West Pico PLUS) and visualized in ImageQuant LAS 4000 mini Biomolecular Imager (GE Healthcare).

### 5.10 Recombinant expression in yeast

Heterologous expression of *Pf*PolK was performed in *Saccharomyces cerevisiae* W303-1A strain. Cells were routinely cultured in liquid or solid YPD medium (2% dextrose, 2% peptone, 1% yeast extract) or liquid/solid Synthetic Defined medium (SD) without the addition of uracil and with 2% dextrose (Bergman, 2001). Yeasts were transformed with either the empty vector (p416-GPD) or with p416-GPD-PfNF54_070015200 (hereafter referred to as p416-*Pf*PolK, i.e., cloned with the *Plasmodium Pf*PolK gene optimized by Genscript for expression in yeast). Yeast expression vectors were transformed into yeasts by the lithium acetate method (Gietz & Woods, 2002). Transformed yeasts were routinely cultured in an SD medium and collected at the early stationary phase for enzymatic assays.

### 5.11 Farnesol and geranylgeraniol kinase activity assays

The recombinant PolK was assayed following the method of Valentin *et al*., for phytol kinase assays (Valentin *et al*, 2006). For this, yeast crude extracts transformed with p416-*Pf*PolK or the empty vector (control) were employed. Yeasts were cultivated until the stationary phase in SD plus dextrose medium and then cells were disrupted by glass beads (0.5 mm Ø) (Avramia & Amariei, 2022). Unbroken cells were discarded by centrifugation at 900 x *g* for 1 min and protein was adjusted to 50 mg/ml with 100 mM Tris/HCl pH 7,4. The reaction was performed in 1.5 mL microtubes by incubating approximately 40 mg of yeast protein with 4 mM MgCl_2_, 800 µM CTP, 10 mM sodium orthovanadate, 0.05% CHAPS and 2 µCi [^3^H] FOH or [^3^H] GGOH. [^3^H] prenol was vacuum-dried as it is commercially distributed in ethanol. The volume was adjusted to 100 µL with 100 mM Tris/HCl pH 7,4 and the reaction was initiated by adding the yeast extract. In some assays, drugs were also added to the reaction or the addition of CTP was omitted as controls. After 30 min of incubation at 37 °C, the reaction was stopped by adding 500 µL of n-butanol saturated in water. The mixture was vortexed, and centrifuged at 12.000 x *g* for 10 min and the organic phase was dried under vacuum. The residue was suspended in 10 µL of n-butanol saturated in water and chromatographed on silica 60 plates (20×20 cm, Merck). Plates were developed for 7-10 cm with isopropyl alcohol/ammonia (32%)/water (6:3:1 by volume). FOH / GGOH standards and the respective phosphates and pyrophosphates were run on the same plate to identify the reaction products and substrates. Standards were visualized with iodine vapor. Finally, the plates were treated with EN3HANCE (Perkin Elmer) and exposed to autoradiography for several days at −70°C. The contrast and brightness of autoradiography scans was adjusted for clarity.

## Author Contributions

MC and IBV contributed to conceptualization, formal analysis, investigation, methodology and writing. AH, TK, AF, MPA, MR, contributed to the investigation, methodology, and formal analysis. AMK, LI, TK and AH also contributed to writing – review & editing. AMK and LI also contributed to project administration, funding acquisition, supervision. TK also contributed to data analyses from the bioinformatics and molecular modelling parts.

## Acknowledgments

We thank Prof. Moritz Treeck (The Francis Crick Institute, London, United Kingdom) for providing the Cre-LoxP *P. falciparum* NF54 strain, and Prof. Xavier Fernández-Busquets and Yunuen Avalos Padilla for providing the anti-ferredoxin-NADP reductase antibody and help in IFA experiments. We also thank the Blood Center of Sírio Libanês Hospital (São Paulo, Brazil), for the gift of erythrocytes and the CSC-Finland for the generous computational resources.

## SUPPORTING INFORMATION

**Figure S1.**
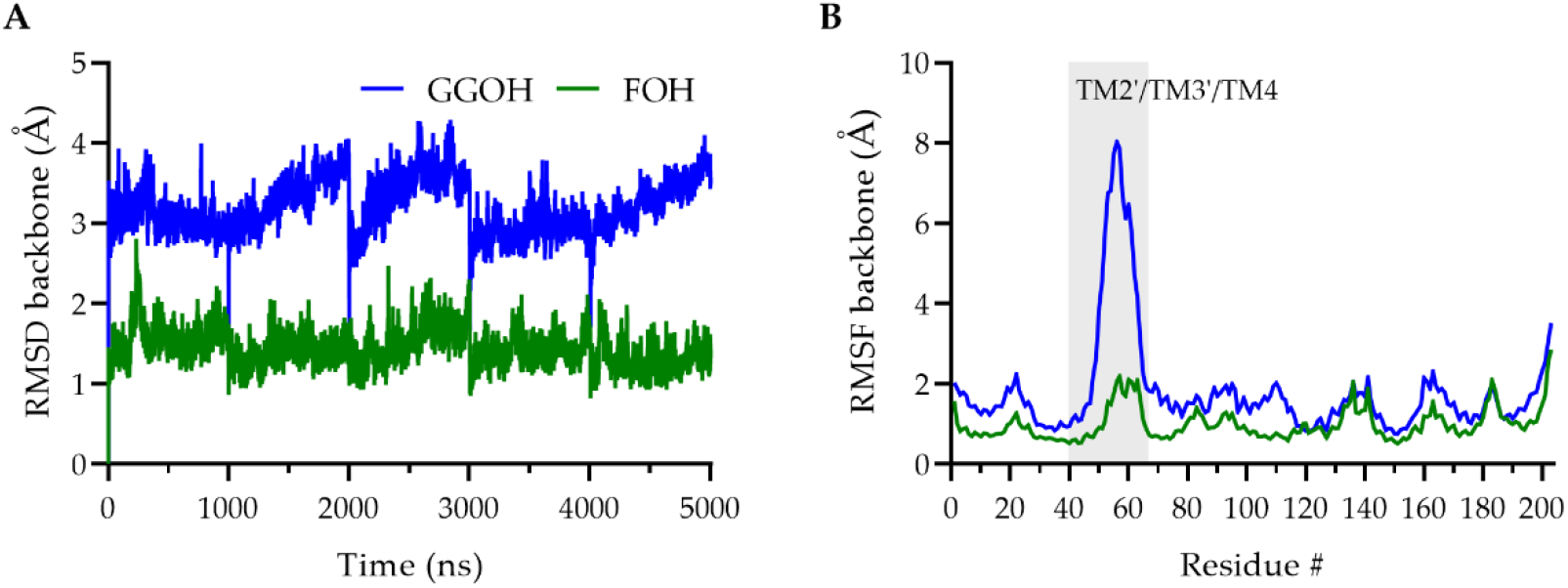
A) Root mean square deviation (RMSD) values of the protein backbone were used to monitor simulation equilibration and protein changes along the trajectory time (merged 5×1 µs). B) Root mean square fluctuation (RMSF) by residues, calculated using the initial MD frame as a reference and compared between ligand-bound, highlighting the TM2’-TM4 region (the intracellular portion) which displays a unique unfolding in the GGOH simulations.

**Figure S2.**
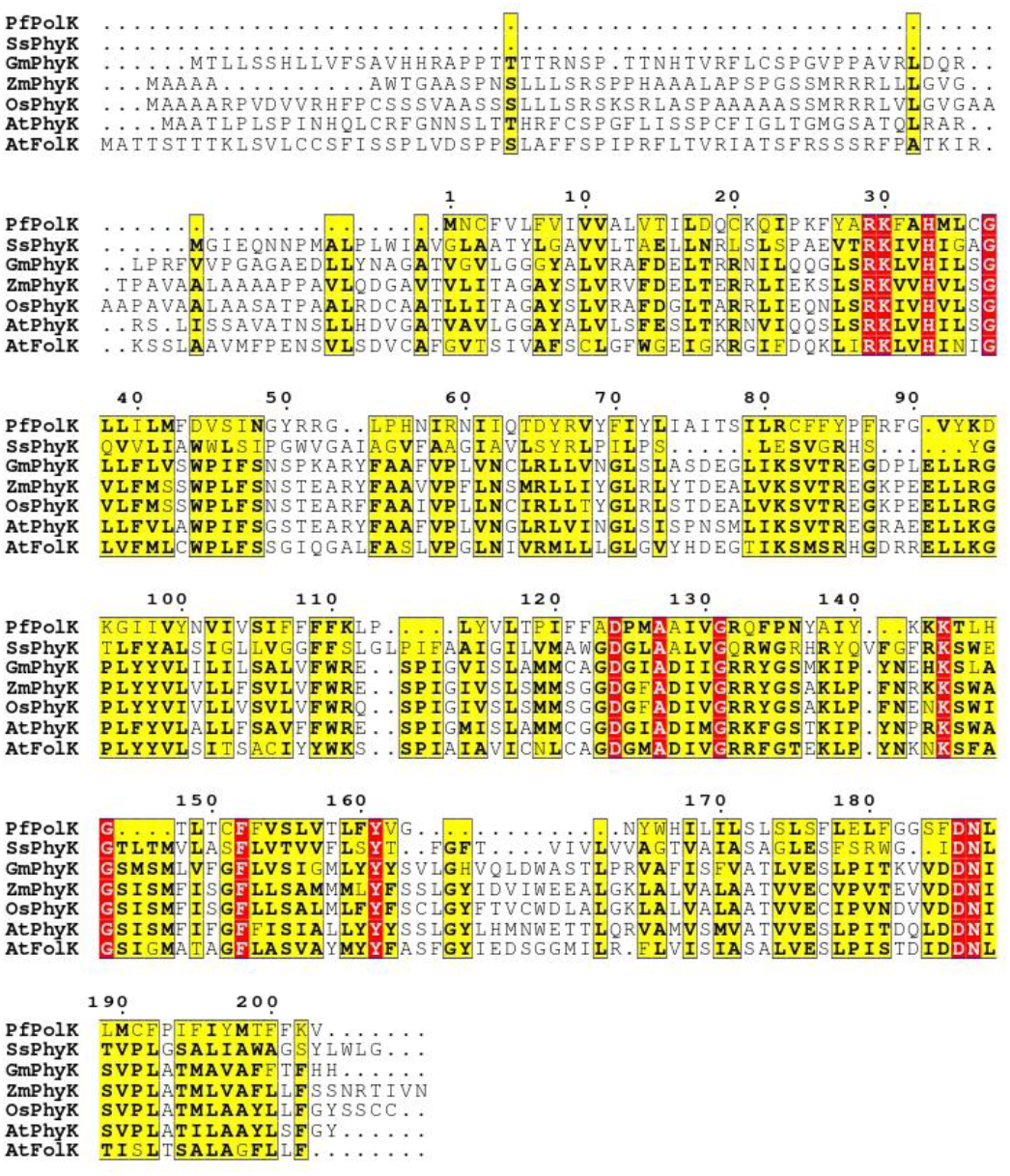
Sequence alignment. ClustalW Multiple alignment of PolK candidate amino acid sequence, predicted in *P. falciparum* NF54 strain against sequences with prenol kinase prediction in Uniprot database. PolK: prenol kinase, PhyK: phytol *kinase*, FolK: farnesol kinase. PfPolK (*P. falciparum* NF54, D0VEH1), SsPhyK (*Synechocystis sp*., P74653), GmPhyK (*Glycine max*, Q2N2K1), ZmPhyK (*Zea mays*, Q2N2K4), OsPhyK (*Oryza sativa*, Q7XR51), AtPhyK (*Arabidopsis thaliana*, Q9LZ76), AtFolK (*Arabidopsis thaliana*, Q67ZM7). Yellow indicates similarity and red indicates identity.

**Figure S3.**
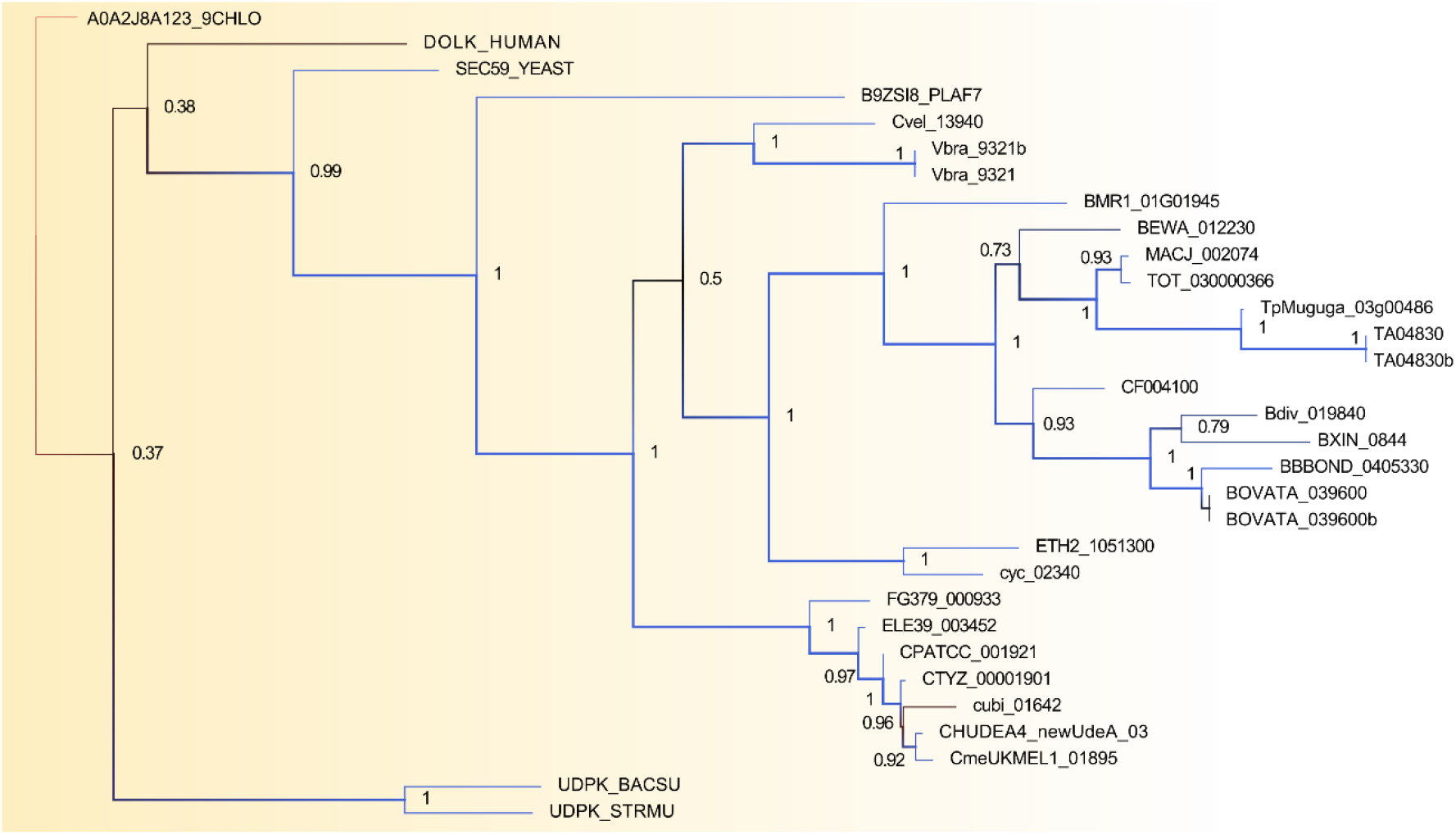
Phylogenetic analysis of the retrieved representative prenol binding proteins. Inset of the overall phylogenetic dendrogram of potential prenol kinases generated using maximum likelihood method (see methods). Branch support values (Bayes posterior probability) are displayed as numbers for the most relevant clade separation, as well as colours (from the highest scores, in blue, to the lowest values, in red) and thickness of the branches. Organisms and genes from Opisthokont group and some extra outliers are highlighted.

**Figure S4.**
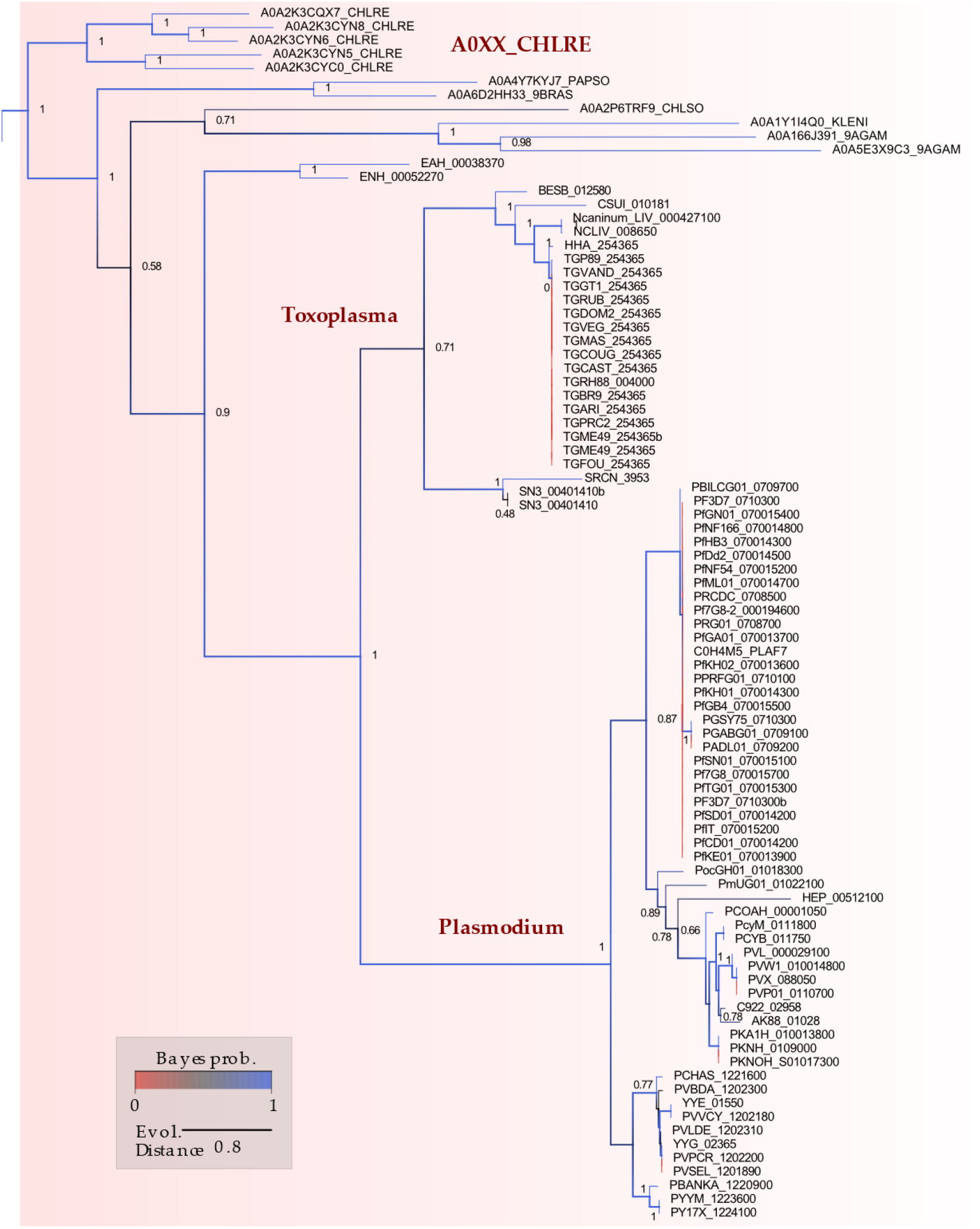
Phylogenetic analysis of dolichol kinase. Inset of the overall phylogenetic dendrogram of potential prenol kinases generated using maximum likelihood method (see methods). Branch support values (Bayes posterior probability) are displayed as numbers for the most relevant clade separation, as well as colours (from the highest scores, in blue, to the lowest values, in red) and thickness of the branches. Apicomplexa clades and the *C. reinhardtii* (A0XX_CHLRE, where A0XX is a generic label for all the *C. reinhardtii*’s taxa) are highlighted.

**Figure S5.**
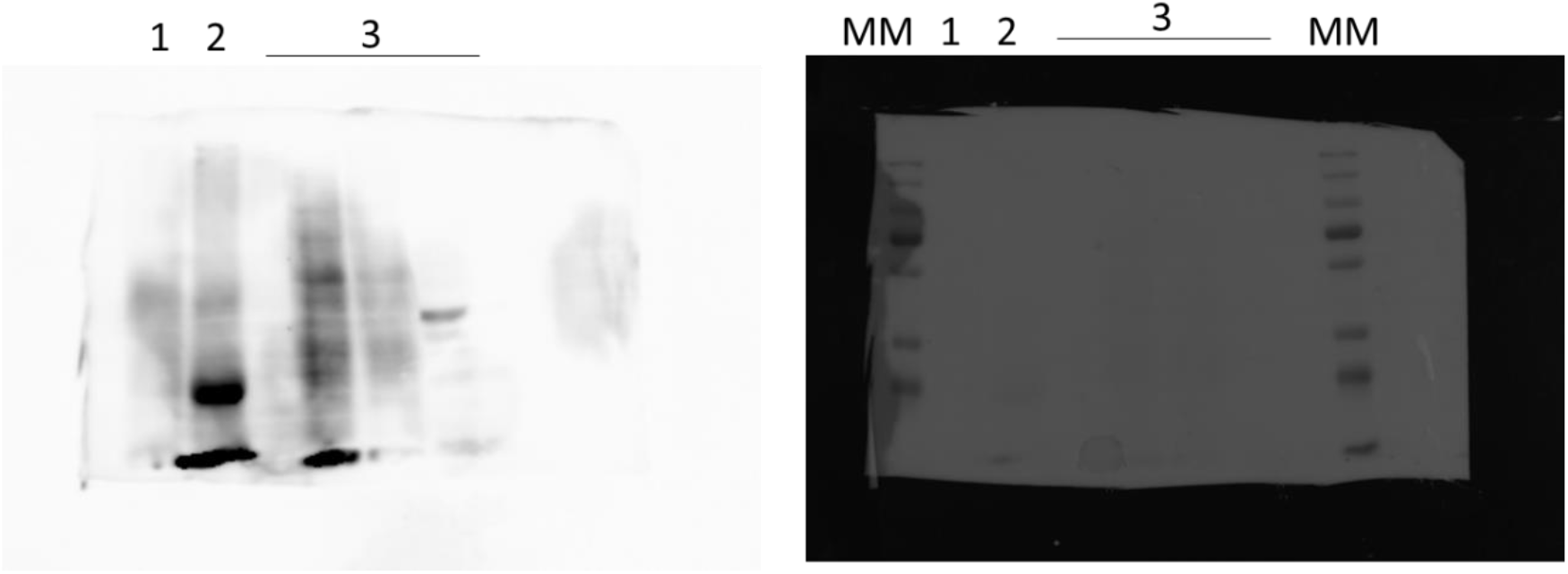
Original images of Western blot analysis. Photographs of Western blot of transgenic parasites (left) and the respective nitrocellulose membrane (right) with the protein ladder (MM). Western blot was performed to analyse the HA-tagged *Pf*PolK of parasites in which *Pf*PolK was excised (Lane 1, parasites exposed to rapamycin) or preserved (Lane 2, parasites exposed to DMSO). The rest of the lanes (group of lanes 3 and onwards) correspond to experiments not related to this article.

